# Visual perceptual learning generalizes to untrained effectors

**DOI:** 10.1101/2020.09.10.291674

**Authors:** Asmara Awada, Shahab Bakhtiari, Christopher C. Pack

## Abstract

Visual perceptual learning (VPL) is an improvement in visual function following training. Although the practical utility of VPL was once thought to be limited by its specificity to the precise stimuli used during training, more recent work has shown that such specificity can be overcome with appropriate training protocols. In contrast, relatively little is known about the extent to which VPL exhibits motor specificity. Indeed, previous studies have shown that training paradigms that require one type of response (e.g., a button press) do not necessarily transfer to those that require a different response (e.g., a mouse movement). In this work, we have examined the effector specificity of VPL by training observers on tasks that maintain the same visual stimuli and task structure, but that require different effectors to indicate the response. We find that, in these conditions, VPL transfers fully between manual and oculomotor responses. These results are consistent with the idea that VPL entails the learning of a decision rule that can generalize across effectors.

## Introduction

Visual Perceptual Learning (VPL) is a long-lasting improvement in the visual system’s ability to detect, to discriminate or to identify visual stimuli following training or experience. For subjects with normal visual acuity, VPL can shed light on fundamental processes such as perceptual development (E. Gibson & Pick, 2003; E. J. Gibson, 1969) and the formation of visual expertise (Appelbaum & Erickson, 2018; DeLoss, Watanabe, & Andersen, 2015; Deveau, Ozer, & Seitz, 2014; Laamerad, Guitton, & Pack, 2020; Reingold & Sheridan, 2011). There is also evidence that VPL can improve outcomes for visual rehabilitation in ageing and clinical populations (Campana, Camilleri, Pavan, Veronese, & Lo Giudice, 2014; DeLoss et al., 2015; Huxlin et al., 2009; Liao, Gichira, Wang, & Chen, 2015; Maniglia, Cottereau, Soler, & Trotter, 2016; Maniglia, Pavan, et al., 2016).

Previous work has shown that healthy populations can be trained to improve their discrimination performance for a wide range of visual features, including orientation (Jehee, Ling, Swisher, van Bergen, & Tong, 2012b; Wang et al., 2016; Xiong, Zhang, & Yu, 2016), contrast (Cong, Wang, Yu, & Zhang, 2016; Yu, Zhang, Qiu, & Fang, 2016), motion (Liang, Zhou, Fahle, & Liu, 2015; Zhang & Yang, 2014) and speed (Yehezkel, Sterkin, Lev, & Polat, 2015). However, most studies report that such learning is highly *specific* to the trained task and the composition of the visual stimulus. Following subtle changes in features such as the stimulus’s location (Hung & Seitz, 2014), orientation (Jehee et al., 2012b) or even the eye of training (Batson, Beer, Seitz, & Watanabe, 2011), the improvement is lost and has to be relearned.

These findings on the specificity of perceptual learning stand in contrast to studies of motor learning, which typically report that the benefits of training are not specific to the trained effector (Albano & Marrero, 1995; Christiansen, Larsen, Grey, Nielsen, & Lundbye-Jensen, 2017; Elliott & Roy, 1981; Imamizu & Shimojo, 1995; Krakauer, Mazzoni, Ghazizadeh, Ravindran, & Shadmehr, 2006; Laszlo, Baguley, & Bairstow, 1970; Modroño et al., 2020; Oosawa, Iwasaki, Suzuki, Tanabe, & Sugawara, 2019; Parlow & Kinsbourne, 1989; Perez et al., 2007; Plow & Carey, 2012; Ruddy & Carson, 2013; Schulze, Lüders, & Jäncke, 2002). For instance, practicing a motor task with one hand leads to improved reaction times that transfer to the other hand, with little or no need for further learning (Gordon, Forssberg, & Iwasaki, 1994; Laszlo et al., 1970; Morton, Lang, & Bastian, 2001). A similar lack of specificity is even observed across very different effectors as, for example, training on an eye movement task improves performance on a hand movement task (Modroño et al., 2020).

Given these results, one might expect the benefits of VPL to transfer to different effectors. That is, the benefits of VPL acquired in a task that requires a response with one effector should persist when a different effector is used. This possibility is seldom tested in VPL experiments, which usually involve a consistent motor response across training and testing. The issue is important, because a pathological specificity of motor responses would severely limit the practical utility of VPL.

Previous studies of motor specificity in VPL have yielded mixed results. While there is some evidence that the benefits of VPL can transfer across effectors, these effects appear to be limited to conditions in which the stimuli and motor responses are the same. Specifically, Grzeczkowski et al. (2019) found that improvements in a VPL task that required a response with one hand persisted when subjects were asked to switch to the other hand. However, in other tasks that entailed a change in the nature of the response (i.e. from a button press to a mouse movement), the benefits of VPL were lost and there was no transfer across effectors (Green, Kattner, Siegel, Kersten, & Schrater, 2015; Grzeczkowski, Cretenoud, Herzog, & Mast, 2017; Grzeczkowski, Cretenoud, Mast, & Herzog, 2019).

These findings can potentially be reconciled under the hypothesis that the specificity of VPL is a property of the decision rule (Green et al., 2015), but not of the motor response *per se* (Szumska, van der Lubbe, Grzeczkowski, & Herzog, 2016). In this case, training on a task that required a binary decision (e.g., right vs. left) would yield benefits that transferred to any effector, but such learning would not transfer to a task that required a continuous readout (e.g., moving a mouse), even if the same effector was used (Grzeczkowski et al., 2019). The finding of inter-manual transfer in VPL (Grzeczkowski et al., 2019) is consistent with this idea, but it might be regarded as a special case, since the movements required for each hand are the same.

In this work, we have tested this hypothesis more generally, using a paradigm in which the stimuli and decision rules were identical across tasks that required responses with very different effectors. Observers were trained on a motion discrimination task that required a binary decision (left vs. right), with the motor response being either a saccade (Experiment 1) or a button press (Experiment 2). Training consistently led to improvements in psychophysical performance (VPL), after which the required motor response was changed. In all observers, we found a full transfer of learning between saccades and manual responses, even though the corresponding effectors are distinct functionally and anatomically. We propose that VPL is not necessarily specific to the motor response and that a perceptual decision rule, once learned, can be flexibly attached to different motor responses as needed.

## Methods

### Observers and Apparatus

Twelve observers with normal or corrected-to-normal vision participated in this study (2 male observers, 10 female observers; 18-25 years). All observers were naïve to the purpose of the study and to visual psychophysics. Observers gave written, informed consent prior to their participation and the study was approved by the Ethics Committee of the Montreal Neurological Institute and Hospital (NEU-06-033). The experiment was halted by the COVID-19 pandemic, but informative data were successfully collected and analyzed from 10 observers who completed the study and 2 who completed almost all of the protocol. As shown below, the results were highly consistent across observers and across tasks.

Observers sat in a normally lit room 40 cm from the monitor, and their heads were stabilized with a chin rest and a forehead bar. Stimuli were generated through the psychophysics toolbox *Psychtoolbox* on MATLAB (Brainard, 1997) and were presented on a 27-inch BenQ monitor (1680 x 1050 pixels, 60 Hz frame rate). Eye position and movements were recorded using the EyeLink 1000 eye tracker (SR Research). Stimuli were viewed binocularly.

### Motion Direction Discrimination Task

#### Stimulus

The stimulus used in this experiment was a translating drifting grating composed of a Gabor patch with a spatial frequency of 1 cycle/degree and a temporal frequency of 6 cycles/second (Figure 1a). The size of the Gabor patch (2 standard deviations of the Gaussian envelope) was 5 degrees, and the stimulus was placed in the upper right quadrant of the visual field at an eccentricity of 5 degrees. This stimulus targets lower-level cortical areas with a high degree of specificity in VPL (Bakhtiari, Awada, & Pack, 2020; Fiorentini & Berardi, 1980; Fiorentini & Berardi, 1981; Hubel & Wiesel, 1959, 1962; Liu & Pack, 2017). Background luminance was 76.48 cd/m^2^, and the contrast was adjusted on each trial according to the staircase procedure described below.

**Figure 1.**
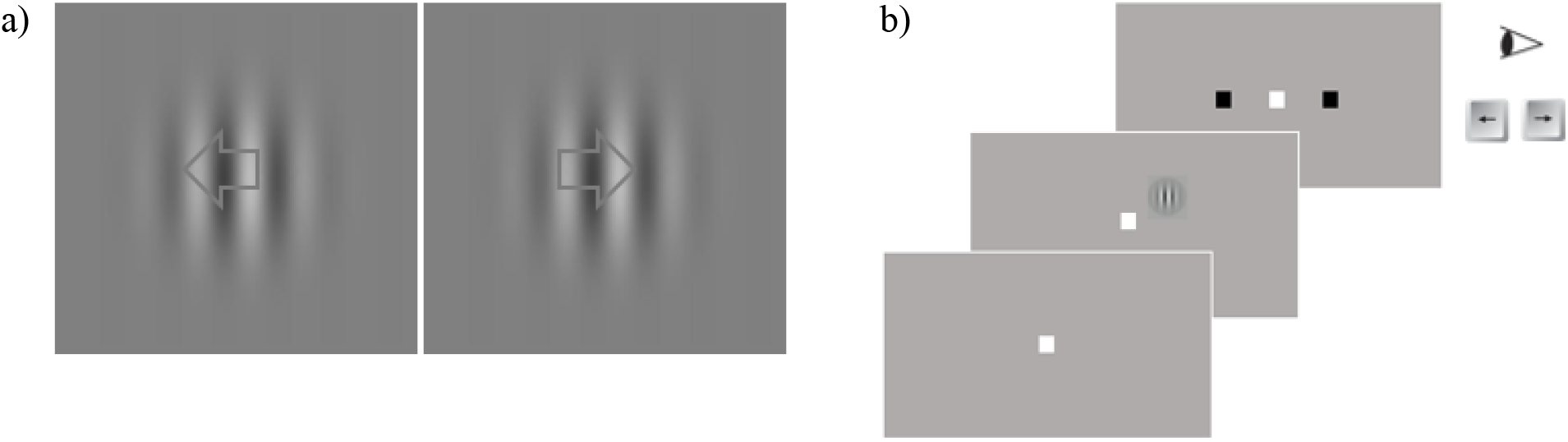
(a) Motion Direction Discrimination Stimulus – Gabor patch drifting to the left or to the right (spatial frequency: 1 cycle/degree, temporal frequency: 6Hz); (b) On each trial, the observer had to fixate on the fixation square for 500ms before the stimulus appeared for four frames (66.7ms), after which two saccade targets appeared, and the observer had to report the direction of the motion with a manual response (keyboard) or a saccade. Each block was composed of 125 trials. One training session was composed of 4 blocks.

#### Task

The experimental task followed a two-alternative-forced-choice paradigm, in which the observer reported the direction of the motion of the stimulus (left or right) through either a manual response (button press on a keyboard) or a saccade.

Each trial started with a central fixation point that the observer had to fixate for 500 ms before the stimulus appeared on the screen for four frames (stimulus duration = 66.7 ms). For trials in which the required response was a saccade, two targets then appeared 5 degrees to the left and right side of the fixation point, at which point the observer had to report the direction of the motion of the stimulus by making a saccade that landed within 1 degree of the target (Figure 1b). For trials in which the required response was a button press, the same two targets appeared, but observers had to press the left or right arrow key on the keyboard. After the response was made, the next trial started.

The direction of motion of the Gabor patch was chosen randomly on each trial to be either right or left. Eye position was tracked throughout the trial and was required to be within 1 degree of the fixation point. If observers broke fixation, the trial was paused until fixation was restored. The starting contrast for the drifting grating was 50%, and the contrast for each subsequent trial was set using a standard 2-down-1-up adaptive staircase procedure (Leek, 2001). Observers were compensated at the rate of 1.2 cents (Canadian) per correct response.

#### Training Paradigm

Each experiment was comprised of two phases. The first phase consisted of one session per day for 7-10 days, and the second phase consisted of one session per day for 5 days. Each session entailed 4 blocks of 125 trials each; the total duration of each session was approximately 30 minutes.

Experiment 1 included 6 observers. In the first phase of Experiment 1, the observers reported the perceived direction of motion with a saccade, and in the second phase, they reported the direction of motion with a manual response.

In the first phase of Experiment 2, the observers reported the direction of the motion with a manual response. In the second phase, the motor response was changed to a saccade. This experiment included a total of 6 observers, 4 of whom successfully completed all phases of the study. Two additional observers successfully completed both phases, with the exception of the last 2-3 sessions, which were halted because of the COVID-19 pandemic.

At the beginning of each phase in both experiments, the experimenter described the task and response required. Observers were unaware of the change in motor response until the second phase, when they were given new instructions about the required motor response.

#### Threshold Measurement and Statistical Analysis

Contrast thresholds were computed using the 2-down 1-up staircase procedure described above, which resulted in an 83% convergence level. Stimulus contrasts at the last six reversals for each training block were averaged, and the threshold for each training session was computed as the median threshold for the 4 blocks run per session.

To quantify generalization to a different motor output, three threshold measurements were computed (Figure 2): the baseline threshold, the training threshold, and the transfer threshold. The baseline threshold represented the threshold during the first training session. The training threshold was computed as the average threshold during the last 5 sessions of phase 1. The transfer threshold was computed as the average threshold during the sessions in phase 2. A standard two-tailed t-test comparison was performed to compare the three threshold values.

**Figure 2.**
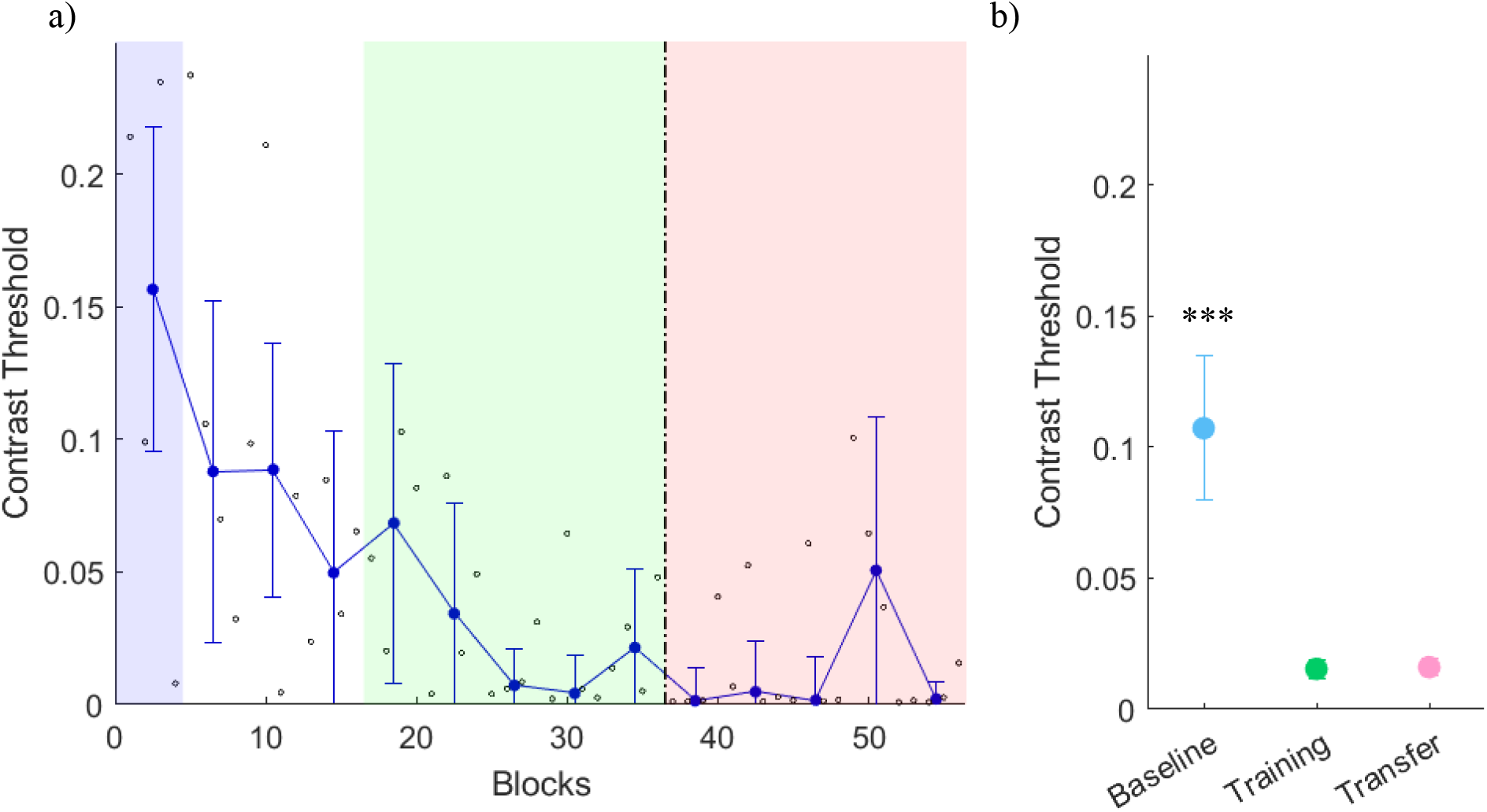
(a) Sample learning curve for one example observer in Experiment 1. The dashed vertical line represents a change in the experimental phase. In phase one (left of the dashed line), the observer reported the direction of the motion with a saccade. In phase two (right of the dashed line), the observer reported the direction of the motion with a manual response (keyboard). Small black dots represent the contrast threshold for each block (125 trials). Blue dots represent the contrast threshold for each training session (median threshold for 4 blocks). Error bars show the standard deviation from the mean contrast threshold for each training session. Shaded regions represent the time periods for each threshold measurement (see (b)); (b) Average thresholds at baseline, during the first phase of training (training) and the second phase of training (transfer) for the six observers in Experiment 1. Baseline threshold is the threshold during the first training session (blue). Training threshold is the threshold during the five last training sessions of phase 1 (green). Transfer threshold is the threshold during the five days of training of phase 2 (pink). Baseline threshold is significantly different from both first-phase training (t(5) = 4.58, *p* =0.006) and second-phase transfer (t(5) = 4.26, *p* = 0.008) thresholds, which are not significantly different from each other (t(5) = −0.186, *p* = 0.86). Error bars show standard deviation from the mean contrast threshold across observers. *** *p* < 0.05.

## Results

We sought to assess whether training with a motion stimulus that shows high levels of sensory specificity would also show motor specificity.

### Experiment 1

In this experiment, we trained 6 observers on a simple motion direction discrimination task with saccades and evaluated whether the improvement would transfer to a manual response (keyboard). Figure 2a shows a sample learning curve for a single subject, with the dashed line indicating the transition from the first experimental phase to the second. If this observer exhibited motor specificity, we would expect to see an abrupt increase in the contrast threshold after the transition, but there was in fact little discernible change (Figure 2a). Indeed, quantifying the results of the six observers shows that the contrast threshold in the second phase was not significantly different from that observed at the end of the first phase (t(5) = −0.186, *p* = 0.86; Figure 2b). Both thresholds were significantly different from the baseline threshold taken on the first session (first phase: t(5) = 4.58, *p* = 0.006; second phase: t(5) = 4.26, *p* = 0.008), indicating that the learning that occurred during the first phase transferred to the second phase. Results for each observer are shown separately in Supplementary Figure 1. Overall, these results indicate that visual perceptual learning transfers when the readout is changed from a saccade to a manual response.

This conclusion is further supported by a comparison of the staircase pattern within different blocks of trials, averaged across observers. As shown in Figure 3a, the staircase during the first (baseline) block shows a slow decline in contrast levels, as observers gradually improved on the task (black line). The last block before the transition from phase 1 to phase 2 shows a rapid decline in contrast, as observers became proficient at the task (purple line). Crucially, the first block after the transition is nearly identical to the last block before the transition (green line), indicating that the proficiency obtained during training transferred almost perfectly to the new motor readout.

**Figure 3.**
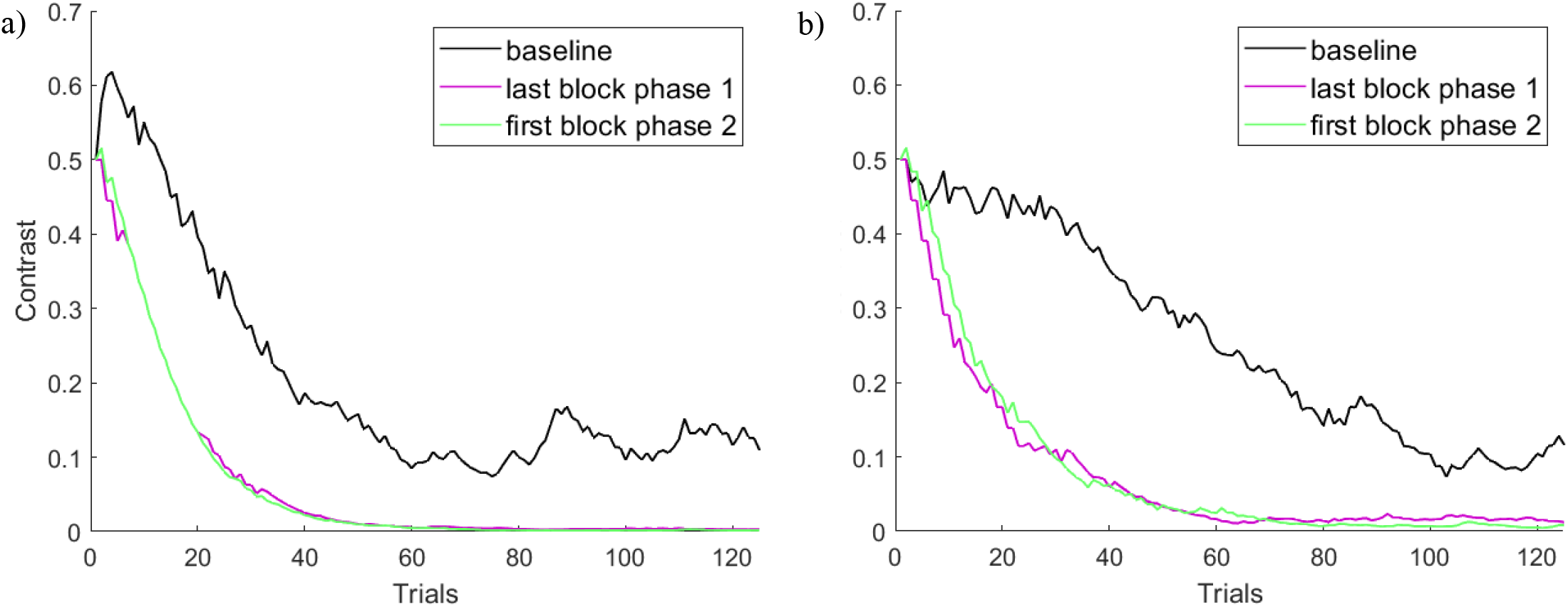
(a) Experiment 1: Staircase pattern at baseline (first block of training with saccade), the last block of training with saccade (phase 1) and the first block of training with a manual response (phase 2). The y-axis represents the average contrast at each trial for the 6 observers; (b) Experiment 2: Staircase pattern at baseline (first block of training with a manual response), the last block of training with a manual response (phase 1) and the first block of training with saccade (phase 2). The y-axis represents the average contrast at each trial for the 6 observers who completed the study.

### Experiment 2

In this experiment, we evaluated whether transfer occurred when changing the motor response from a manual response to a saccade. Six observers completed the training and transition components of the study, although two of these observers were unable to complete the last two or three sessions of phase 2 because of the COVID-19 pandemic. Figure 4a shows a sample learning curve for a single observer, with the dashed line representing the transition from the manual motor response to the saccade. Again, it is clear that the contrast threshold changed very little in the transition from the first to the second phase. As in Experiment 1, we found for the population of observers that thresholds did not change significantly across the transition (t(5) = 0.84, *p* = 0.4389; Figure 4b), and that baseline thresholds differed significantly from those obtained at the end of the first phase (t(5) = 3.74, *p* = 0.0135) and the beginning of the second phase (t(5) = 3.40, *p* = 0.0193). Similar results were obtained when we excluded the two observers who did not complete the full protocol (baseline threshold vs. first-phase training (t(3) = 2.47, *p* = 0.0905; baseline vs. second-phase transfer (t(3) = 2.51, *p* = 0.0869; first-phase vs. second-phase thresholds (t(3) = 2.34, *p* = 0.102). Results for each individual observer are shown in Supplementary Figure 2.

**Figure 4.**
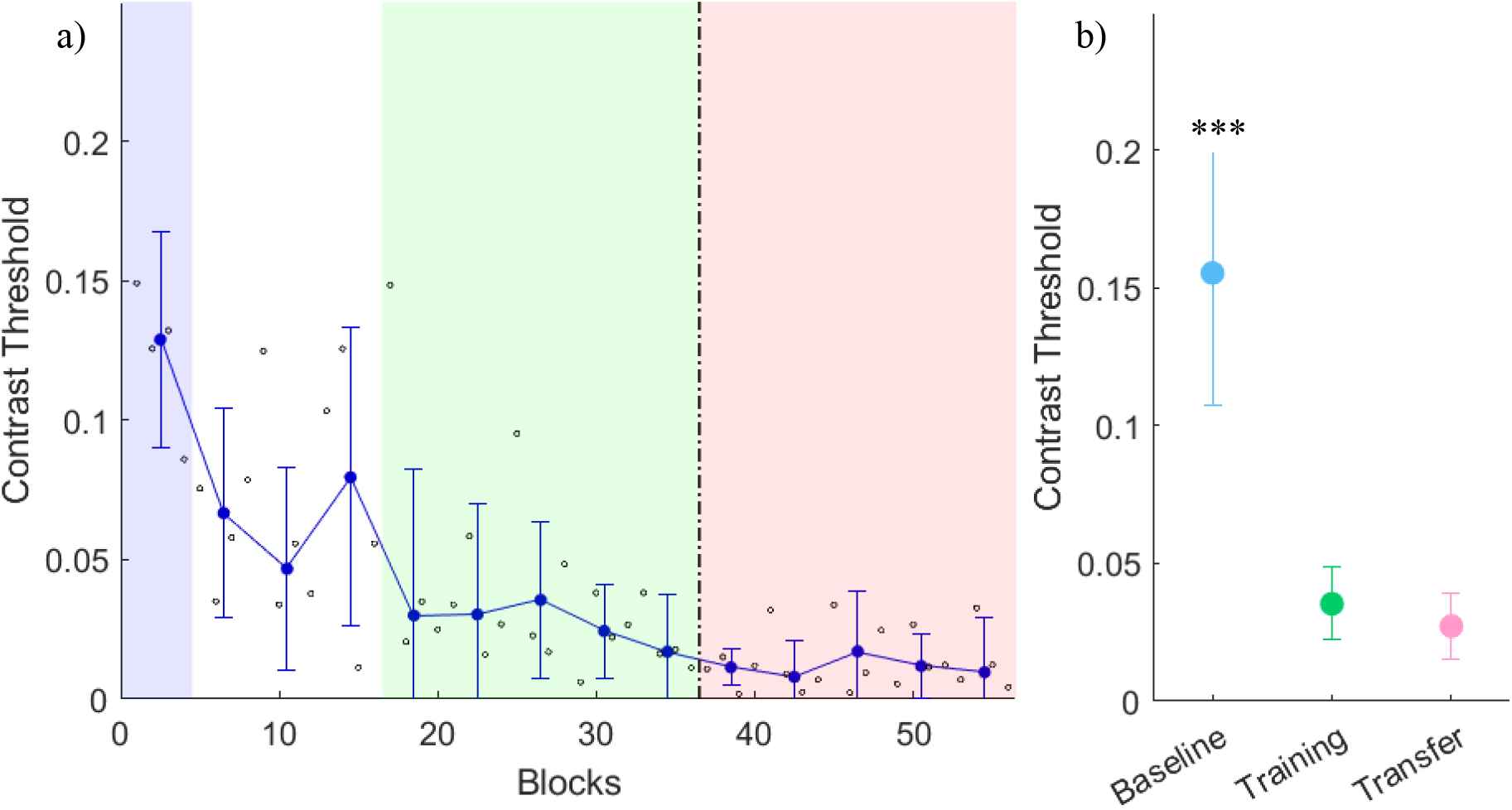
(a) Sample learning curve for one observer in Experiment 2. The vertical dashed line represents the change in experimental phase. In phase one (left of the dashed line), the observer reported the direction of the motion with a manual response (keyboard). In phase two (right of the dashed line), the observer reported the direction of the motion with a saccade. Small black dots represent the contrast threshold for each block (125 trials). Blue dots represent the contrast threshold for each training session/day (median threshold for 4 blocks). Error bars show the standard deviation from the mean contrast threshold for each training session/day. Shaded regions represent the time periods for each threshold measurement (see (b)); (b) Average thresholds at baseline (blue), during the first phase of training (green) and the second phase of training (pink) for six observers. Baseline threshold is significantly different from both first-phase training (t(5) = 3.74, *p* = 0.0135; Figure 3c) and second-phase transfer (t(5) = 3.40, *p* = 0.0193; Figure 3c) thresholds, which are not significantly different from each other (t(5) = 0.84, *p* = 0.4389; Figure 3c). Error bars show standard deviation from the mean contrast threshold across observers. *** *p* < 0.05.

As in Experiment 1, a closer examination of the staircase patterns shows that the progression of contrast values was very similar between the last block of phase 1 and the first block of phase 2, with neither block being similar to baseline (Figure 3b). Overall, these results indicate that significant transfer can occur when changing the motor response from a manual response to a saccade.

## Discussion

Visual perceptual learning (VPL) represents a type of adult cortical plasticity that has significant theoretical and practical implications. Indeed, exploring VPL and its underlying mechanisms can shed light on essential brain functions in the adult visual system and can be used to develop training strategies for those seeking visual expertise or visual rehabilitation. Research in the field has thoroughly focused on exploring the sensory aspect of VPL and has identified a hallmark sensory specificity that might limit its practical use (Batson et al., 2011; Dosher & Lu, 2017; Hung & Seitz, 2014; Jehee, Ling, Swisher, van Bergen, & Tong, 2012a). However, recent advances have highlighted a variety of ways in which this sensory specificity can be overcome (Ahissar & Hochstein, 1997; Green et al., 2015; Talluri, Hung, Seitz, & Seriès, 2015).

At the same time, if VPL is to be of practical utility, it should not be limited to the effector used in the training protocol. Yet to date, little is known about the motoric aspects of VPL. In this work, we propose that the hallmark specificity of VPL does not necessarily extend to motor outputs. Indeed, we show that following training with a stimulus that often yields VPL with high levels of sensory specificity, the improved performance nevertheless transfers to different motor responses. While our sample size may not have been sufficient to detect a small cost of transferring VPL across effectors, the large differences in thresholds at baseline and at transfer (Figures 2b and 4b) suggest substantial motor generalization.

### Implications for VPL

Our results have a number of implications for a mechanistic understanding of VPL. First, our findings are in line with the proposal that two separate anatomical pathways exist for object perception and motor responses (Goodale & Milner, 1992). Indeed, our results suggest that the improved perception of a stimulus in VPL occurs independently of the response to that stimulus. Our results are also consistent with most VPL theories that posit that visual processing follows an information processing framework in which perception occurs before any decision making or action occurs (Li, 2016; Marr, 1982; Watanabe & Sasaki, 2015).

Second, our results shed light on the long-standing debate about the brain locus of VPL. A number of studies have shown that VPL can reweight the readout of visual information, emphasizing the cortical regions most suitable for the trained task (Bakhtiari et al., 2020; Chang, Mevorach, Kourtzi, & Welchman, 2014; Chen, Cai, Zhou, Thompson, & Fang, 2016; Chowdhury & DeAngelis, 2008; Liu & Pack, 2017; Walsh, Ashbridge, & Cowey, 1998). In our experiments, the most suitable brain regions were presumably low-level cortical areas, which optimally encode grating stimuli of the kind used in our training protocol (Bakhtiari, Awada, & Pack, 2020; Fiorentini & Berardi, 1980; Fiorentini & Berardi, 1981; Hubel & Wiesel, 1959, 1962; Liu & Pack, 2017). Anatomical and physiological evidence suggests that direct connections between these visual areas and motor regions are weak and that these connections exhibit a strong preference for saccades over limb movements (Levy, Schluppeck, Heeger, & Glimcher, 2007; Strigaro et al., 2015). This suggests an intermediate stage of processing between sensory and motor regions, which is capable of encoding perceptual decision rules independently of the motor response.

One possibility is the parietal lobe, more specifically the posterior parietal cortex (PPC), which has been shown to be responsible for sensorimotor transformations in visually-guided behaviors in non-human primates (Andersen & Buneo, 2002; Andersen & Cui, 2009; Andersen, Essick, & Siegel, 1987; Andersen, Snyder, Bradley, & Xing, 1997). Indeed, Law and Gold demonstrated that training non-human primates on a motion perception task with saccades resulted in changes in the neural response in the lateral intraparietal area (LIP) of the PPC and not in visual area MT, which is responsible for motion perception (Law & Gold, 2008). There is ample evidence that PPC neurons can respond to both saccades and arm movements, even in specialized areas such as LIP and the parietal reach region (PRR) (Snyder, Batista, & Andersen, 1997). For instance, a recent fMRI study by Levy and associates found only slight preferences for a specific effector in the human equivalents of areas LIP and PRR (Levy et al., 2007). In the same fMRI study, Levy and associates further demonstrated that a greater effector specificity existed in visual and motor areas outside parietal cortex, as early visual areas were activated during saccades and motor areas during reaching movements (Levy et al., 2007). Our results thus support a framework in which VPL reflects a change in the efficiency with which sensorimotor structures in the parietal lobe read out visual information from the occipital cortex (Chen et al., 2016; Law & Gold, 2008).

In this regard, the PPC might be the site at which decision rules – mappings from sensory stimuli to perceptual outputs – reside. Recent evidence suggests that perceptual learning can occur at a ‘conceptual level’ at which abstract rules, rather than specific stimulus mappings, are learnt (Green et al., 2015; Wang et al., 2016). Differences in VPL specificity might thus be attributed to differences in a learned decision “rule”: a rule that encourages flexibility, via a task that exposes the observer to multiple stimulus conditions and responses, will naturally lead to less specificity, and this has been observed experimentally (Bavelier, Achtman, Mani, & Föcker, 2012; Green et al., 2015). Such rules need not be specific to the motor response, and indeed our results suggest that once a decision rule is learnt, it can flexibly be mapped onto untrained motor responses. At the same time, binary decision rules might be regarded as a special case (Szumska, van der Lubbe, Grzeczkowski, & Herzog, 2016), and it remains to be seen whether other types of rules would also generalize to different effectors.

### Implications for vision rehabilitation

Our findings have important implications for the practical utility of VPL, namely for rehabilitation. VPL has been shown to be somewhat successful in recovering some visual function in patients with V1 lesions, although the improvements seem to be limited by a high degree of specificity to the stimulus and its location (Cavanaugh & Huxlin, 2017; Das, Tadin, & Huxlin, 2014; Huxlin et al., 2009; Sahraie et al., 2006). Such specificity can to some degree be overcome with the use of training stimuli that target higher-level visual areas, for which generalization at the neural level is greater (Das et al., 2014).

Even if stimulus specificity could be overcome, motor specificity could sharply limit the utility of VPL in vision rehabilitation, and in this regard, the finding that learning is not necessarily specific to the motor response reinforces its use as a potential therapy. That said, the above considerations on decision rules suggest that rehabilitation protocols should encourage flexibility in mapping stimuli to motor responses. As mentioned previously, VPL appears to be highly sensitive to the decision rules embodied by the training task (Green et al., 2015); specificity often arises because observers learn to link individual stimuli to individual response. Thus, it is likely useful to avoid protocols that require a continuum of visual stimuli and motor outputs (Achtman, Green, & Bavelier, 2008).

## Acknowledgements

This work was supported by a grant from NSERC to C.C.P. We thank Dr. Michael Herzog for helpful discussions.

