## Supplementary Figures 1 and 2 for "Visual perceptual learning generalizes to untrained effectors"

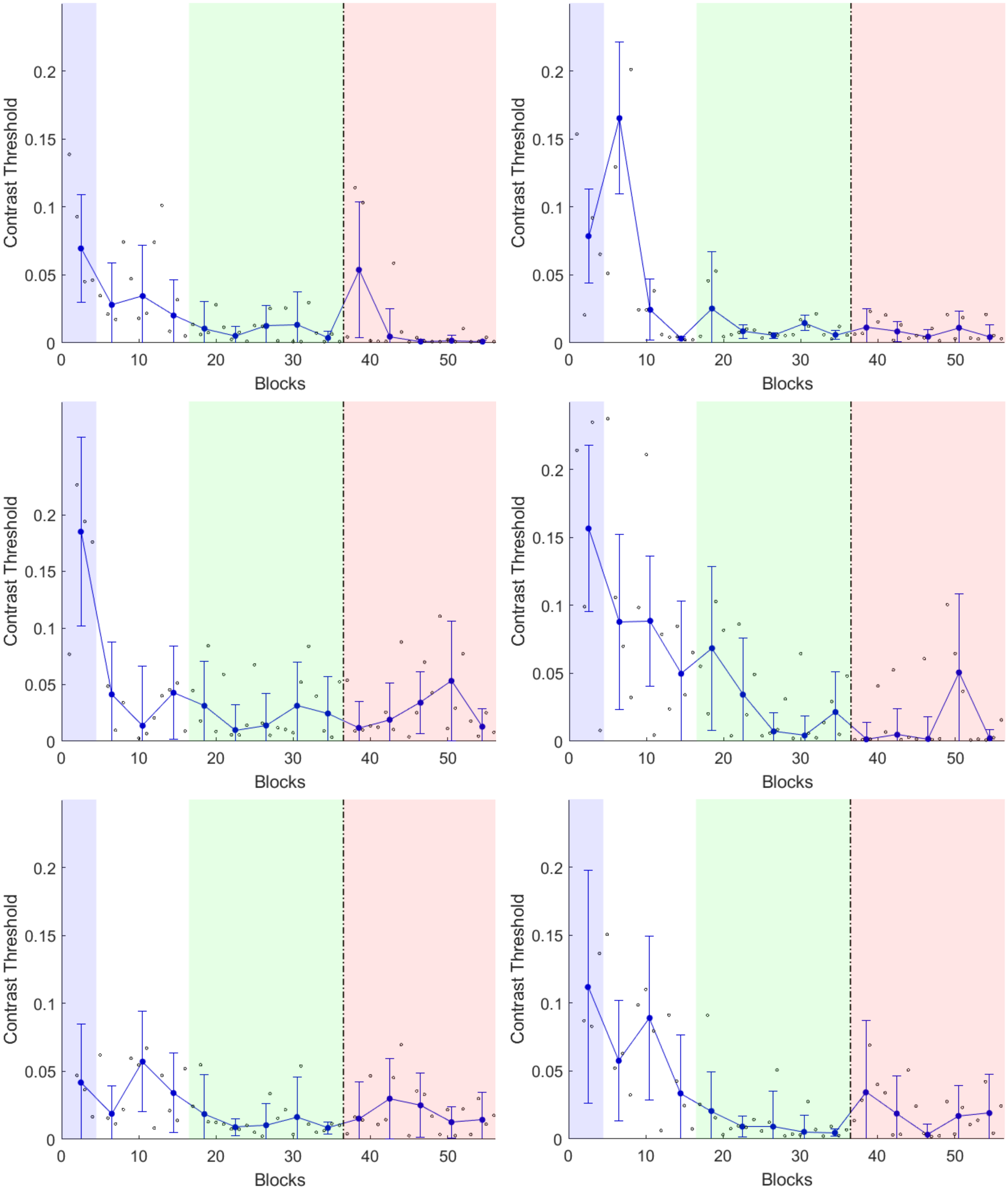

**Figure S1** Learning curves for each observer (n = 6) in Experiment 1. The dashed line represents the change in experimental phase. In phase one (left of the dashed line), the observer reported the direction of the motion with a saccade. In phase two (right of the dashed line), the observer reported the direction of the motion with a manual response (keyboard). Small black dots represent the contrast threshold for each block (125 trials). Blue dots represent the contrast threshold for each training session/day (median threshold for 4 blocks). Error bars show the standard deviation from the mean contrast threshold for each training session/day. Shaded regions represent the time periods for each threshold measurement. Baseline threshold (blue) was computed as the threshold during the first day of training with the saccade. The training threshold was computed as the threshold during the last five days of training with the saccade (green). The transfer threshold was computed as threshold during the five days of training with the manual response (pink).

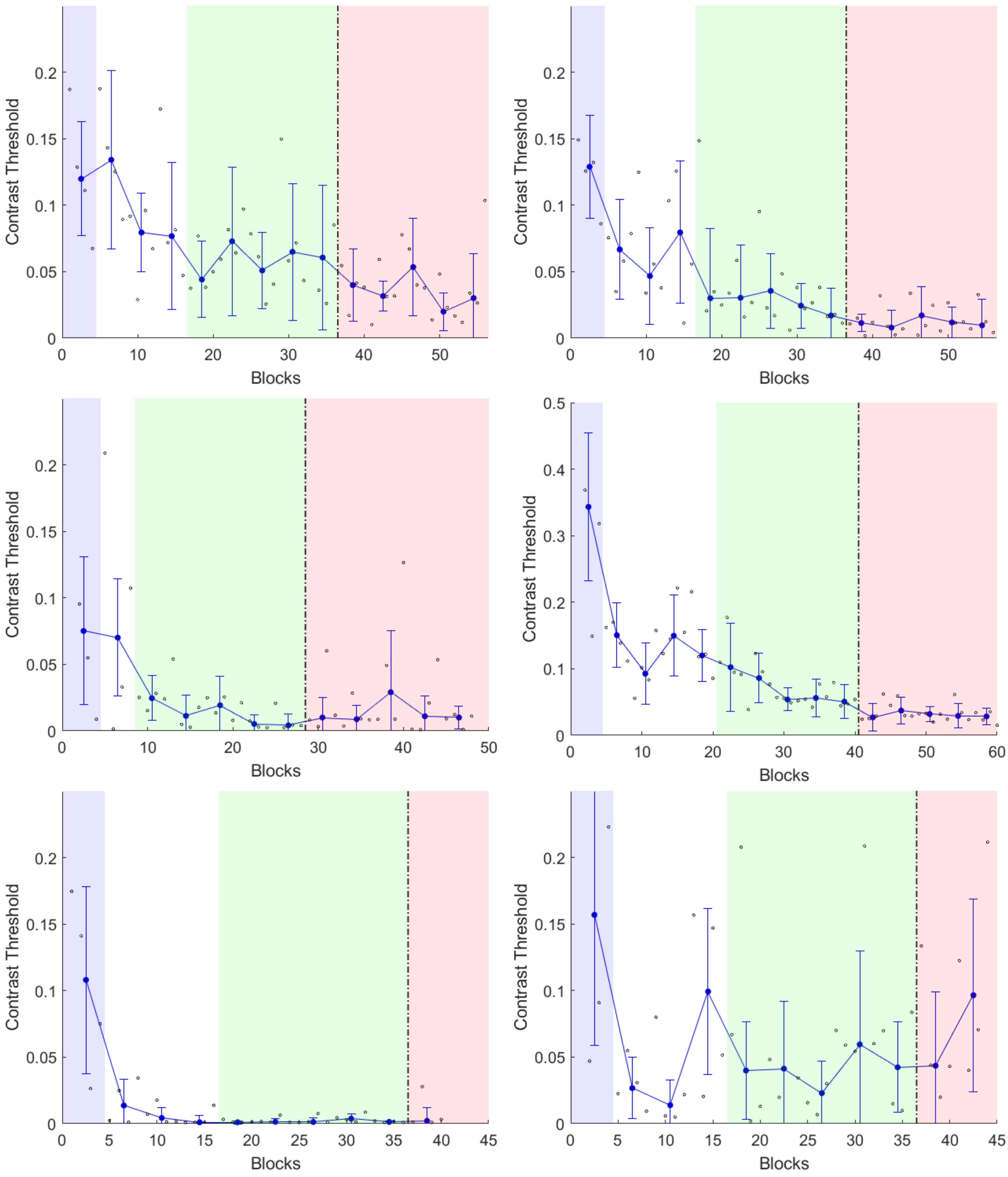

Contrast Threshold
Blocks

**Figure S2** Learning curves for each observer (n = 6) in Experiment 2. The dashed line represents the change in experimental phase. In phase one (left of the dashed line), the observer reported the direction of the motion with a manual response (keyboard). In phase two (right of the dashed line), the observer reported the direction of the motion with a saccade. Small black dots represent the contrast threshold for each block (125 trials). Blue dots represent the contrast threshold for each training session/day (median threshold for 4 blocks). Error bars show the standard deviation from the mean contrast threshold for each training session/day. Shaded regions represent the time periods for each threshold measurement. Baseline threshold (blue) was computed as the threshold during the first day of training with the manual response. The training threshold was computed as the threshold during the last five days of training with the manual response (green). The transfer threshold was computed as threshold during the five days of training with the saccade (pink). Due to the COVID-19 pandemic, two observers (bottom row) were unable to complete the five days of training in phase 2 in Experiment 2.
